# Sudden unexpected death in epilepsy is prevented by blocking postictal hypoxia

**DOI:** 10.1101/2022.03.25.485818

**Authors:** Antis G. George, Jordan S. Farrell, Roberto Colangeli, Alexandra K. Wall, Renaud C. Gom, Mitchell T. Kesler, Cristiane L de la Hoz, Tefani Perera, Jong M. Rho, Deborah Kurrasch, G. Campbell Teskey

## Abstract

Epilepsy is at times a fatal disease. Sudden unexpected death in epilepsy (SUDEP) is the leading cause of mortality in people with intractable epilepsy and is defined by exclusion; non-accidental, non-toxicologic, and non-anatomic causes of death. While SUDEP often follows a bilateral tonic-clonic seizure, the mechanisms that ultimately lead to terminal apnea and then asystole remain elusive and there is a lack preventative treatments. Based on the observation that discrete seizures lead to local vasoconstriction, resulting in hypoperfusion, hypoxia and behavioural disturbances in the forebrain (Farrell et al., 2016), we reasoned that similar mechanisms may play a role in SUDEP when seizures invade the brainstem. Here we tested this neurovascular-based hypothesis of SUDEP in awake non-anesthetized mice by pharmacologically preventing seizure-induced vasoconstriction, with cyclooxygenase-2 or L-type calcium channel antagonists. In both acute and chronic mouse models of SUDEP, ibuprofen and nicardipine extended life. We also examined the potential role of spreading depolarization in the acute model of SUDEP. These data provide a proof of principle for the neurovascular hypothesis of SUDEP and the use of currently available treatments to prevent it.

## Introduction

Sudden unexpected death in epilepsy (SUDEP) is the non-accidental, non-toxicologic, and non-anatomic cause of death in people with epilepsy. SUDEP accounts for up to 18% of deaths in epilepsy, which exceeds the expected rate in the general population by nearly 24 times (Nashef et al., 2007). It is the leading cause of mortality in young adults with drug-resistant epilepsy with an annual incidence of 1 per 1000 (Hesdorffer et al., 2011; Thurman et al., 2014). Amongst neurological disorders, SUDEP is second only to stroke in the number of potential years of life lost (Ficker et al., 1998; Tomson et al., 2008; Thurman et al., 2014). There are currently no effective treatments to prevent SUDEP in people with intractable epilepsy.

The brain mechanisms underlying SUDEP are largely unknown but *postictal* respiratory dysfunction led to terminal apnea, which preceded cessation of heartbeat (asystole), in the largest retrospective review of SUDEP and near-SUDEP cases; the MORTality in Epilepsy Monitoring Units Study (MORTEMUS) (Ryvlin et al., 2013). Before preventative strategies can be implemented, the pathophysiology of SUDEP must be elucidated. Multiple potential causes of SUDEP have emerged from clinical and experimental studies including cardiac dysfunction (Friedman et al., 2018; Szurhaj et al., 2021), automimic dysfunction (Lee and Devinksy, 2005), arousal dysfunction (Petrucci et al., 2020), and intrinsic airway obstruction (Stewart et al., 2017). In recent years spreading depolarization (SD) has been studied in SUDEP pathophysiology (Aiba and Noebels, 2015; Loonen et al., 2019; Jansen et al., 2019). SD is a self-propagating wave of depolarization that silences neuronal networks. Although SD can be evoked experimentally by high potassium, tetanic neuronal stimulation, or mechanical damage, it can also arise during limited energy substrate availability such as brain hypoxia (Dreier, 2011).

Postictal vasoconstriction-induced hypoperfusion and hypoxia is found to occur in local brain regions involved in seizures (Farrell et al., 2016; Gaxiola-Valdez et al., 2017; Li et al., 2019; Liu et al., 2020; Farrell et al., 2020) and is long and severe enough to result in the postictal state and cause behavioural dysfunction (Farrell et al., 2017). Moreover, two drug targets have been identified that effectively prevent postictal hypoperfusion/hypoxia and behavioural dysfunction without inhibiting seizures; cyclooxygenase 2 (COX-2) and L-type calcium channels (LTCCs) (Farrell et al., 2016). Extending the work of Farrell and colleagues, Tran et al., using 2-photon microvasculature and Ca^++^ imaging in awake mice reported robust vasoconstriction of cortical penetrating arterioles at the onset of an induced seizure which lasted over an hour (Tran et al., 2020). The observed vasoconstriction corresponded to a rise in vascular smooth muscle cell Ca^++^ which persisted beyond the onset of the seizure. Further, inhibition of COX-2 with ibuprofen and LTCC’s with nicardipine prevented this severe vasoconstriction (Farrell et al., 2016, Tran et al., 2020). As cases of SUDEP are strongly linked to an ictal event (Walczak et al., 2001; Ryvlin et al., 2013), postictal-induced local hypoperfusion/hypoxia in the brainstem provides a neurovascular-based hypothesis for the neurobiological mechanisms underlying SUDEP and there are readily available drugs to test this hypothesis (Farrell et al., 2016; Farrell et al., 2020).

In the present study, we utilized two seizure models that produce bilateral tonic-clonic seizures followed by premature mortality during the postictal period: acute intrahippocampal kainic acid that results in several bouts of seizures (Puttachary et al., 2015) and *Kcna1*^-/-^ mice that display self-generated seizures (Rho et al., 1999; Glasscock et al., 2007; Simeone et al., 2018). Both models recapitulate the central features observed in the MORTEMUS study as well as postictal hypoperfusion/hypoxia. While the COX-2 inhibitor, ibuprofen, and the LTCC antagonist, nicardipine, ameliorate postictal hypoxia through fundamentally different mechanisms, both drugs significantly extended life. Using the acute model in non-anesthetized mice, we also observed that brainstem SD was a consequence rather than a cause of seizure-induced brainstem hypoxia.

## Results

### Acute hippocampal kainic acid administration in mice replicates human SUDEP

We first determined whether mortality following acute intrahippocampal kainate infusion in mice replicated the sequence of electrographic seizure, terminal apnea followed by asystole observed in clinical SUDEP. We infused kainic acid into the hippocampus of male C57BL/6J (n=4) and *Kcna1*^-/-^ (n=4) mice and recorded local field potentials (LFP) and absolute partial pressure of oxygen (pO_2_) in the pre-Bötzinger complex (PBC). The PBC has been studied extensively and is a considered a critical node for central respiratory rhythmogenesis and eupnea (normal breathing) (Ramirez et al., 2012). The PBC is well characterized in mammals, including humans (Schwarzacher et al.,2011). Evidence from clinical SUDEP cases suggest brainstem respiratory involvement and as such the PBC was targeted for our investigation (Ryvlin et al., 2013). We also recorded respiratory rate, and heart rate (**Fig. 1A**). Following a terminal bilateral tonic-clonic seizure which lasted a mean of 27.52 ± 3.37 seconds (**Fig. 1B**), all mice had a precipitous fall in pre-Bötzinger complex pO_2_ (**Fig. 1C**), followed by respiratory distress, presenting as agonal breathing, bradypnea and finally apnea (**Fig.1D**). Asystole then occurred 7.32 ± 1.2 minutes after the terminal apneic event (**Fig. 1E**). Thus, the acute intrahippocampal kainic acid mouse model replicates the essential sequence of events observed in human SUDEP cases (**Fig.1F**) (Ryvlin et al., 2013).

**Fig 1:**
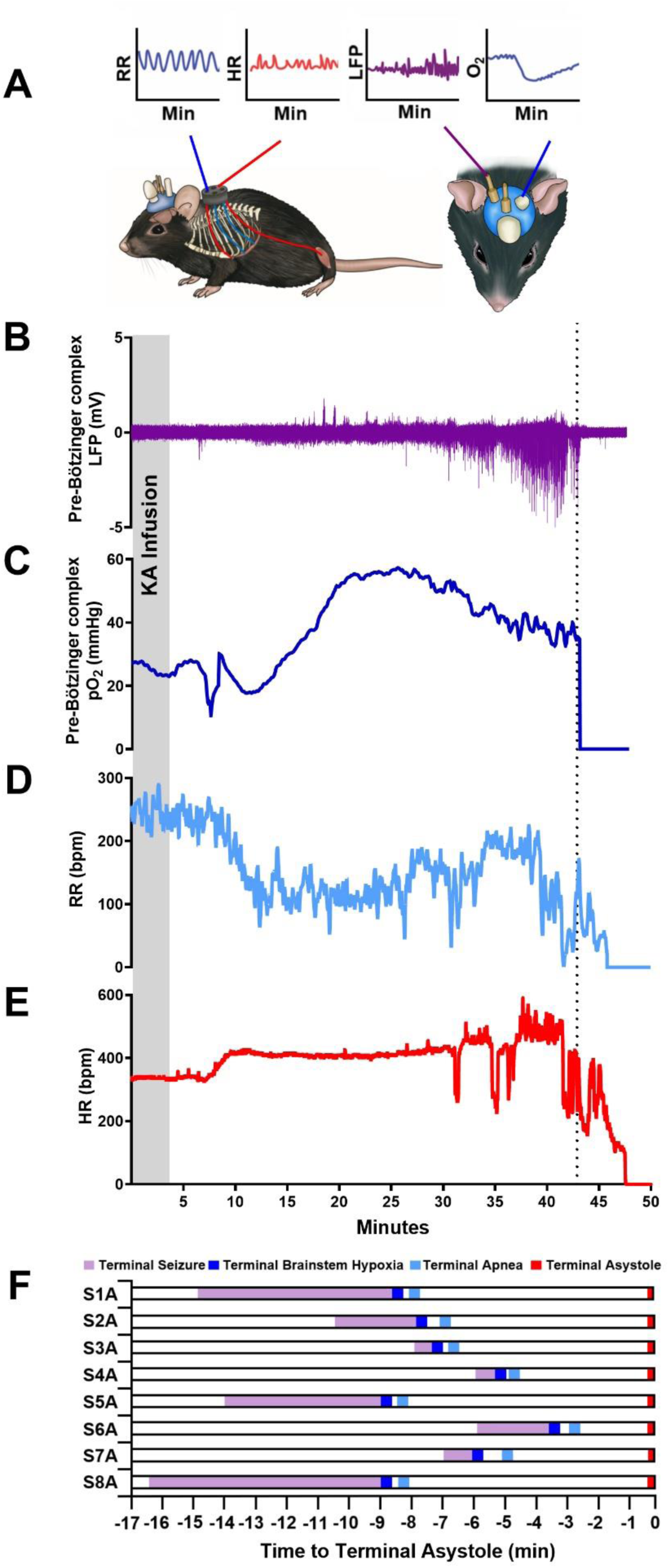
Modelling SUDEP: Respiratory failure precedes asystole in an acute mouse model. Representative profile from a C57BL/6J + kainic acid mouse (no differences were observed between C57BL/6J and *Kcna1*^-/-^ mice) during a terminal seizure. **A**. Model schematic: Electromyograph (EMG) electrode wires record respiratory activity directly from diaphragm and electrodes record cardiac activity. A bipolar electrode in the brainstem records local field potential (LFP) in the pre-Bötzinger complex (PBC) and an oxygen sensing optode in the contralateral PBC records absolute oxygen. **B-E**. Electrographic seizures precede a drop in pO_2_ levels in respiratory nuclei followed by terminal apnea. Epileptiform activity in the PBC indicates seizure propagation to the brainstem. The mouse had a terminal fully generalized seizure lasting 1.48 minutes, immediately followed by a precipitous decline in pO_2_ in the PBC. **D**. Respiration rate (RR) arrested 1.11 minutes after PBC became severely hypoxic (<10mmHg). **E**. Heart rate (HR) continued for 3.24 minutes following cessation of breathing. **F**. Graphical representation of male C57BL/6J (n=4; S1A to 4A) and *Kcna1*^-/-^ (n=4; S5A to 8A) mice highlighting the sequence of events leading to terminal asystole. In all recorded mice a stereotyped sequence of events occurred. All mice displayed a full generalized terminal seizure (purple), the PBC became severely hypoxic (dark blue), resulting in terminal apnea (light blue), however, terminal asystole (red) did not occur until several minutes after terminal apnea.

### Postictal hypoxia precedes terminal apnea and asystole in brainstem PBC and NTS

The neurovascular hypothesis of SUDEP predicts that following electrographic seizures in brainstem respiratory nuclei pO_2_ will drop, which in turn leads to cessation of breathing (Farrell et al., 2017). Thus, we recorded respiration, terminal electrographic seizures and pO_2_ levels at the site of seizure origin, the hippocampus, as well as two brainstem respiratory nuclei in male C57BL/6J (n=6) and *Kcna1*^-/-^ (n=6) mice (**Fig. 2)**. We targeted both the pre-Bötzinger complex, located in the ventral medulla, which is responsible for the generation of rhythmic inspiratory breathing movements (Akins et al., 2017) and the nucleus of the solitary tract (solitary nucleus; NTS) located in the dorsal medulla which is a critical node for cardio-respiratory function (Gasparini et al., 2020). At baseline, before infusion of kainic acid, the pre-Bötzinger complex (35.54 mmHg ± 3.38) and NTS (36.12 mmHg ± 2.36) displayed significantly higher mean pO_2_ levels relative to the hippocampus (20.68 mmHg ± 1.45). (**Fig. 2 D-D*i***). During terminal seizures, epileptiform activity in the hippocampus preceded and terminated before epileptiform activity in both brainstem nuclei (**Fig. 2 A-C**). pO_2_ levels decreased dramatically toward the end of a terminal electrographic seizure. We observed that the pre-Bötzinger and NTS becomes severely hypoxic prior to apnea (2.5 ± 0.54 seconds). **D*i***. Once pO_2_ levels dropped below 3.4 ± 0.28 mmHg in the pre-Bötzinger complex, breathing ceased.

**Figure 2:**
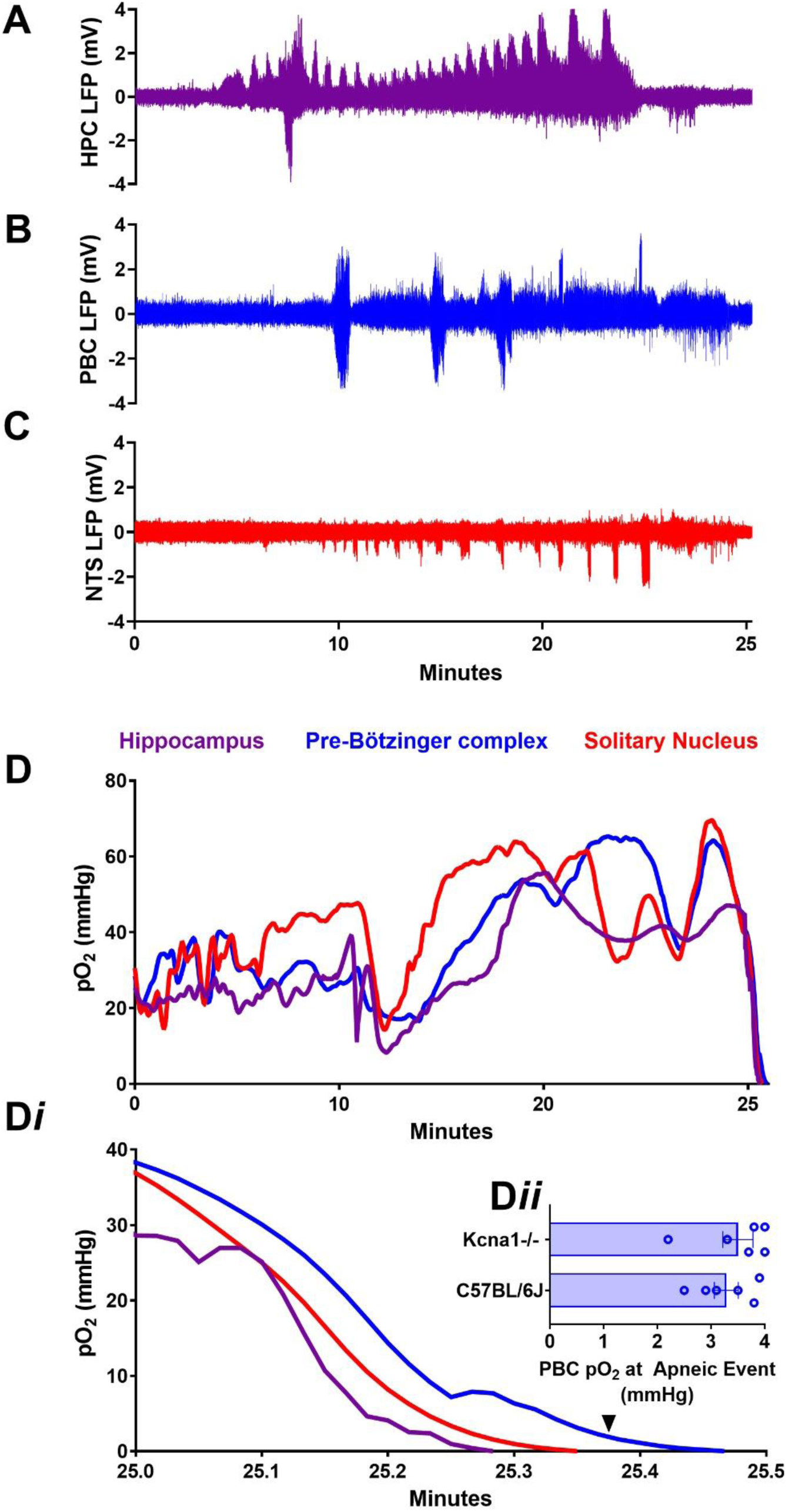
Electrographic seizures precede a drop in pO_2_ levels in respiratory nuclei followed by terminal apnea. Representative profile from a representative mouse during a terminal seizure following kainic acid infusion. **D-D*i***. Baseline pre-Bötzinger complex (PBC; blue line) and Nucleus of the solitary tract (solitary nucleus; NTS; red line) displayed higher pO_2_ levels relative to the hippocampus (HPC; purple line) (35.54 mmHg ± 3.38, 36.12 mmHg ± 2.36, 20.68 mmHg ± 1.45, respectively. ANOVA, F_2,33_=12.02 ^***^p=0.0001). **A-C**. During terminal seizures epileptiform activity in the hippocampus (purple) ended earlier relative to PBC (blue) and NTS (red). **D**. pO_2_ levels decreased dramatically toward the end of a terminal electrographic seizure (0.2 and 4.3 minutes). Both the pre-Bötzinger complex and NTS becomes severely hypoxic prior to apnea. Once pO_2_ levels dropped below 4 mmHg in the pre-Bötzinger complex, breathing ceased (black arrow). During the terminal apneic period, pre-Bötzinger complex and the NTS oxygen ranged between 0.0 and 4.0 mmHg. **D*i***. Expanded inset outlines terminal hypoxia order, hippocampus reaches anoxia first followed by NTS and finally PBC. **D*ii***. Histobars representing PBC pO_2_ at terminal apneic event in male C57BL/6J (n=6) and *Kcna1*^-/-^ (n=6) mice.

### Blocking postictal hypoxia extends life in both males and females

COX-2 inhibitors and LTCC antagonists prevent postictal vasoconstriction/hypoperfusion/hypoxia in the rodent forebrain (Farrell et al., 2016; Tran et al., 2020; Farrell 2021). Thus, we predicted that pre-treatment with a single bolus of the COX-2 inhibitor ibuprofen would extend life until its effective concentration was metabolically eliminated. After intrahippocampal administration of kainic acid, male and female mice treated with vehicle, or a low dose of ibuprofen (15 mg/kg) displayed terminal electrographic seizures in the brainstem and then became terminally apneic and died at asystole at similar time points. However, male, and female mice treated with 50 or 100 mg/kg ibuprofen also displayed electrographic seizures around the same time that the vehicle mice died but brainstem pO_2_ remained within the normoxic range and breathing was not disrupted **Suppl Fig. 1**. After ibuprofen was metabolized to lower levels (2 to 3.3 half-lives; Salama et al., 2016) brainstem seizures led to pO_2_ levels below 4 mmHg, and terminal apnea ensued. Thus, pre-administration of ibuprofen dose-dependently and significantly extended life, independent of a change in seizure duration or frequency (**Fig.3 A-F**).

**Fig 3:**
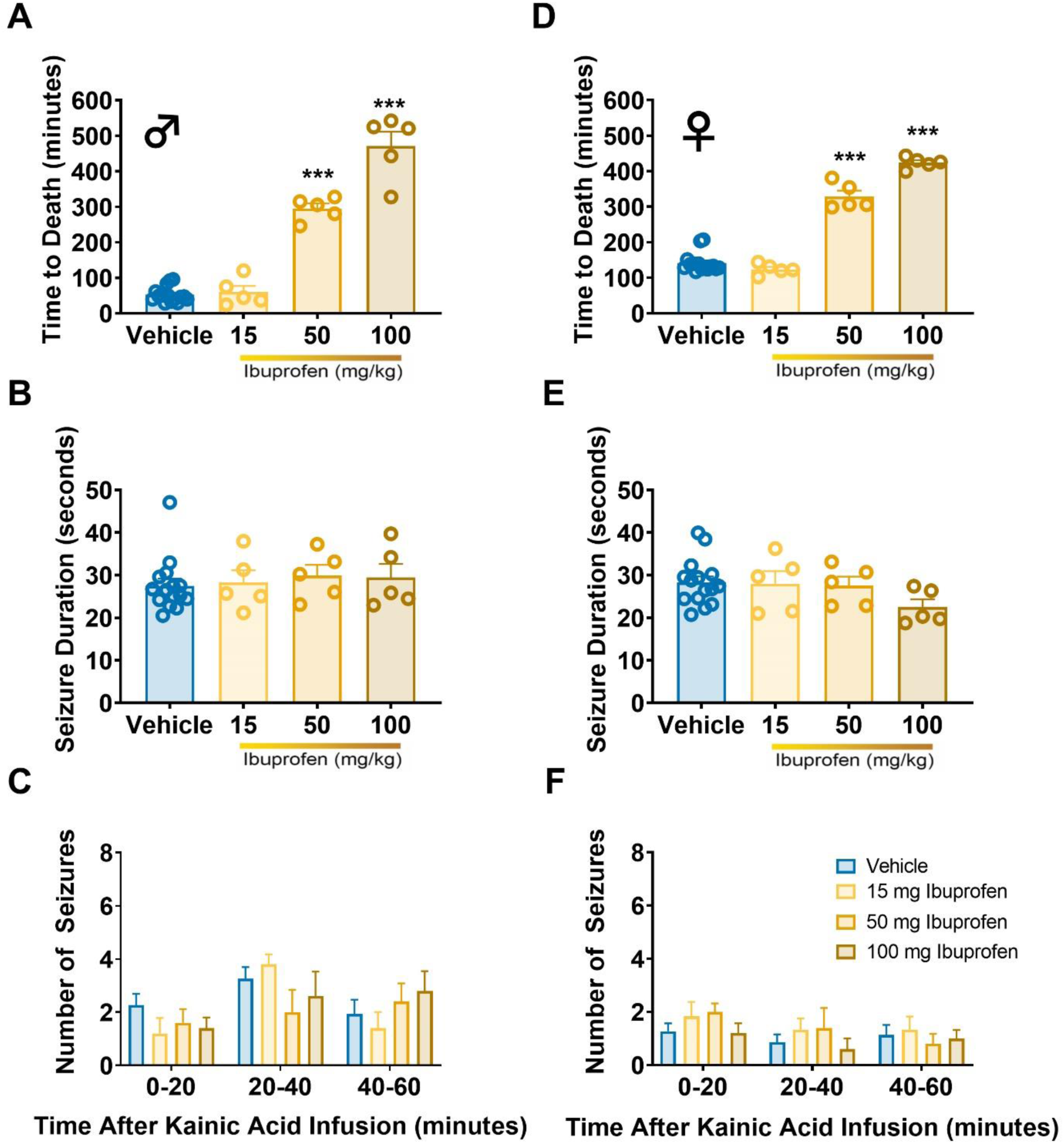
Ibuprofen dose-dependently extends survival time in C57BL/6J + kainic acid mice. **A**. The mean time to death in ibuprofen treated male mice is significantly greater than that for DMSO treated controls (ANOVA, F_3,26_ =143 ^***^p=<0.0001). **D**. Female mice also had a delay in mortality compared to that of DMSO treated controls (ANOVA, F_3,26_=198.1 ^***^p=<0.0001). **B. E. C. F**. There was also no difference in seizure profiles with respect to overall seizure duration (Male: ANOVA, F_3,26_=0.23 p=0.87; Female: ANOVA, F_3,26_=1.52 p=0.23) and number of seizures (Male: 2-Way ANOVA, F_6,78_=1.13 p=0.35; Female: F_6,81_=0.327 p=0.92) in all groups. Histobars represent means ± SEM.

To further test the idea that multiple dosing of ibuprofen could further extend life in the face of repeated seizures we next delivered repeated dosages of ibuprofen to keep plasma levels relatively high. Male C57BL/6J vehicle control mice (n=5) died 61.04 ± 12.30 minutes after kainic acid infusion. However, mice (n=5) that received an initial loading dose of 100 mg/kg ibuprofen followed by 3 successive doses of 50 mg/kg ibuprofen every 2 hours, equivalent to ibuprofen’s half-life (Salama et al., 2016), lived 574.6 ± 16.27 minutes (9.58h) after kainic acid infusion and significantly longer than those that received only a single dose of ibuprofen. Thus, we were able to prevent death while mice had sufficient levels of the COX-2 inhibitor in their system.

The intrahippocampal kainic acid model continues to generate seizures for hours in mice dosed with ibuprofen. In clinical practice, patients actively seizing (i.e. seizures that do not self-terminate), receive pharmacological intervention to terminate seizures, protect brain function and prevent death (Bilington et al., 2016). Here we sought to enhance the therapeutic efficacy of ibuprofen in extending life, by terminating seizures with either diazepam or a combination of diazepam with either ketamine or isoflurane. Diazepam, a benzodiazepine, was chosen as it is considered a first-line agent in the management of acute seizures and status epilepticus (Orchoa and Kilgo, 2016) followed by second-line agents which consist of anesthetics (Mirsattari et al., 2004; Fang and Wang, 2015). Male C57BL/6J (n=5) control mice that did not receive ibuprofen or antiseizure treatment had a median survival of 46.8 minutes after kainic acid infusion. All treatment groups received 100 mg/kg ibuprofen 30 minutes before kainic acid infusion. The median survival time was longer in mice treated with ketamine (n=5) was 4 days post infusion. Mice that received ibuprofen and diazepam had a median survival day of 5. Mice that received ibuprofen, ketamine and diazepam had a median survival of 5 days. Finally, mice that received ibuprofen, diazepam and isoflurane had a median survival of 6 days post infusion of kainic acid (**Fig. 4**). Mice that received polypharmacological intervention to terminate seizures then lived beyond our two-day observation window provide evidence that greater survival can be achieved. All mice did eventually die regardless of treatment arm due to the relatively high dose of kainic acid chosen to model seizure-induced premature mortality.

**Fig. 4.**
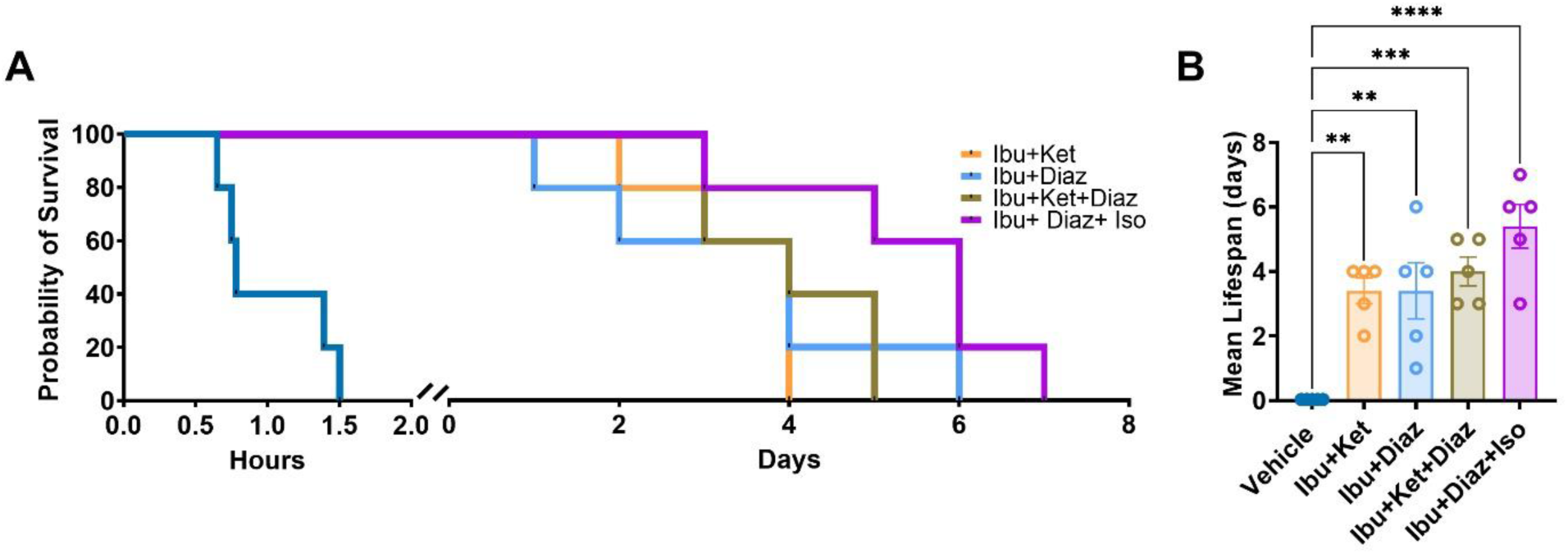
Termination of seizure activity increases survival in ibuprofen treated C57BL/6J mice following intrahippocampal kainic acid administration. **A**. The median survival for vehicle treated mice (dark blue trace; n=5) was 46.8 minutes post kainic acid infusion. All treatment groups received 100 mg/kg ibuprofen 30 minutes before kainic acid infusion. The median survival for mice also treated with ketamine (yellow trace; n=5) was 4 days. The median survival for mice also treated with diazepam (light blue trace; n=5) was 5 days. The median survival for mice also treated with ketamine and diazepam (brown trace; n=5) was 5 days. The median survival of mice also treated with diazepam and isoflurane (purple trace; n=5) was 6 days (p= 0.0001; Mantel-Cox test). **B**. All treatment conditions significantly extended survival (ANOVA, F_4,20_ =12.33 ^****^p=<0.0001).

To test the neurovascular hypothesis of SUDEP more thoroughly we employed the drug, nicardipine, that also prevents seizure-induced vasoconstriction but with a non-overlapping mechanism of action with COX-2 inhibitors. LTCC drugs were previously determined to block postictal hypoxia without inhibiting seizures (Farrell et al., 2016). Nicardipine was chosen because it is routinely given in emergent patients to treat cerebral vasospasm and its ability to cross the blood-brain-barrier effectively (Keyrouz and Diringer, 2007). We administered nicardipine (5 mg/kg) i.p. 30 minutes before acute intrahippocampal kainic acid. Male and female control mice (n=5/sex) died 58.90 ± 12.63 and 129.6 ± 6.182 minutes respectively after kainic acid infusion (**Fig. 5)**. However, male and female mice pre-treated with nicardipine (n=5/sex) lived 143.4 ± 5.6 and 188.4 ± 4.4 minutes respectively after kainic acid infusion. Interestingly, male mice treated with nicardipine had significantly longer seizure duration compared to controls and female nicardipine treated counterparts. The reason for this increase is currently unknown. We show that, life extension by nicardipine, a drug that prevents postictal hypoxia by a different mechanism than COX-2 inhibitors, provides convergent evidence for the neurovascular hypothesis of SUDEP.

**Fig. 5:**
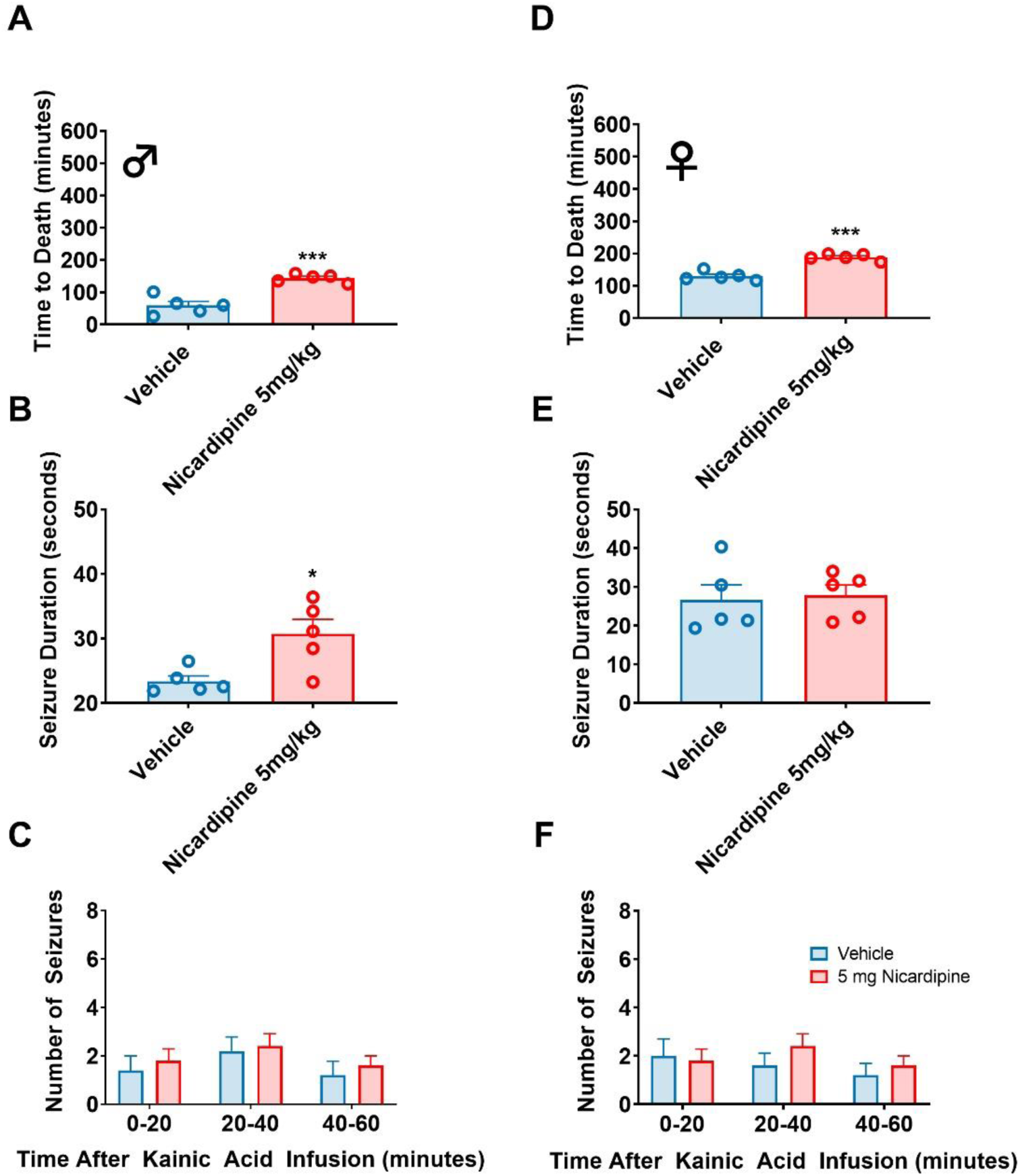
Administering the L-type calcium channel blocker nicardipine extends survival time in C57BL/6J + kainic acid mice. A. The mean time to death in nicardipine-treated male mice is significantly greater than that for DMSO treated controls t8=6.10, ***p=0.0003 and D. female t8=7.74 ****p=<0.0001 C57BL/6J mice. B.C. Male mice treated with nicardipine had a significant increase in overall seizure duration compared to vehicles t8=2.98, *p=0.0175. E.F. There was no difference in overall seizure duration in female mice (t8=0.252, P=0.81) or total number of seizures (Male: 2-Way ANOVA, F2,24=0.023 p=0.98; Female: 2-Way ANOVA, F2,24=0.45 p=0.64). Histobars represent means ± SEM.

### Terminal brainstem spreading depolarization does not precede but follows SUDEP in non-anesthetized mice

Brainstem spreading depolarization (SD) has been reported to be associated with SUDEP pathophysiology in some transgenic mouse models using anesthetic preparations (Aiba and Noebels, 2015; Aiba et al., 2016; Cain et al., 2022). Further, several rodent studies using general anesthesia have reported an association between brain hypoxia and SD (Funke et al., 2009; Richter et al., 2010). To assess whether brainstem hypoxia initiates brainstem SD or vice versa, we performed alternating current (AC) and direct current (DC) electrographic recordings as well as simultaneous oxygen, heart, and respiratory recordings from 5 male C57BL/6J and 5 male *Kcna1* ^*-/ -*^ head-fixed (non-anesthetized) mice in the acute SUDEP model.

Following kainic acid infusion, there was at least one non-fatal electrographic seizure, then a drop in oxygen followed by a negative DC-potential shift and then a recovery in oxygen level. All mice displayed this stereotyped sequence of events leading to death (**Fig. 6**). Our data without the confound of anesthesia clearly indicate that brainstem hypoxia precedes apnea which precedes onset of terminal SD. Baseline pO_2_ in the PBC was 41.6 ± 2.2 mmHg and remained at this value until the primary AD electrographic seizure event. During the primary seizure event severe hypoxia (pO_2_ < 10 mmHg) occurred. The lowest recorded value for non-fatal seizure pO_2_ was 9.2 ± 1.0 mmHg and hypoxia lasted 1.5 ± 0.8 minutes before returning to normoxic levels. During the fatal terminal AC-recorded seizure, severe hypoxia preceded SD by 11.5 ± 2.06 minutes indicating hypoxic SD. Hypoxic SD, which shares many features of SD is hallmark of temporary regional or global hypoperfusion and ischemia in the brain (Czeh et al., 1993; Dreier and Reiffurth, 2015; Zandt et al., 2011). SD occurred 36.7 ± 3.1 seconds before terminal asystole. These data indicate that fatal SD occurs in the brainstem once it becomes terminally hypoxic. The negative DC shift observed in raw LFP recordings occurred after a fatal seizure and long after hypoxia commenced, indicative of anoxic depolarization.

**Fig. 6.**
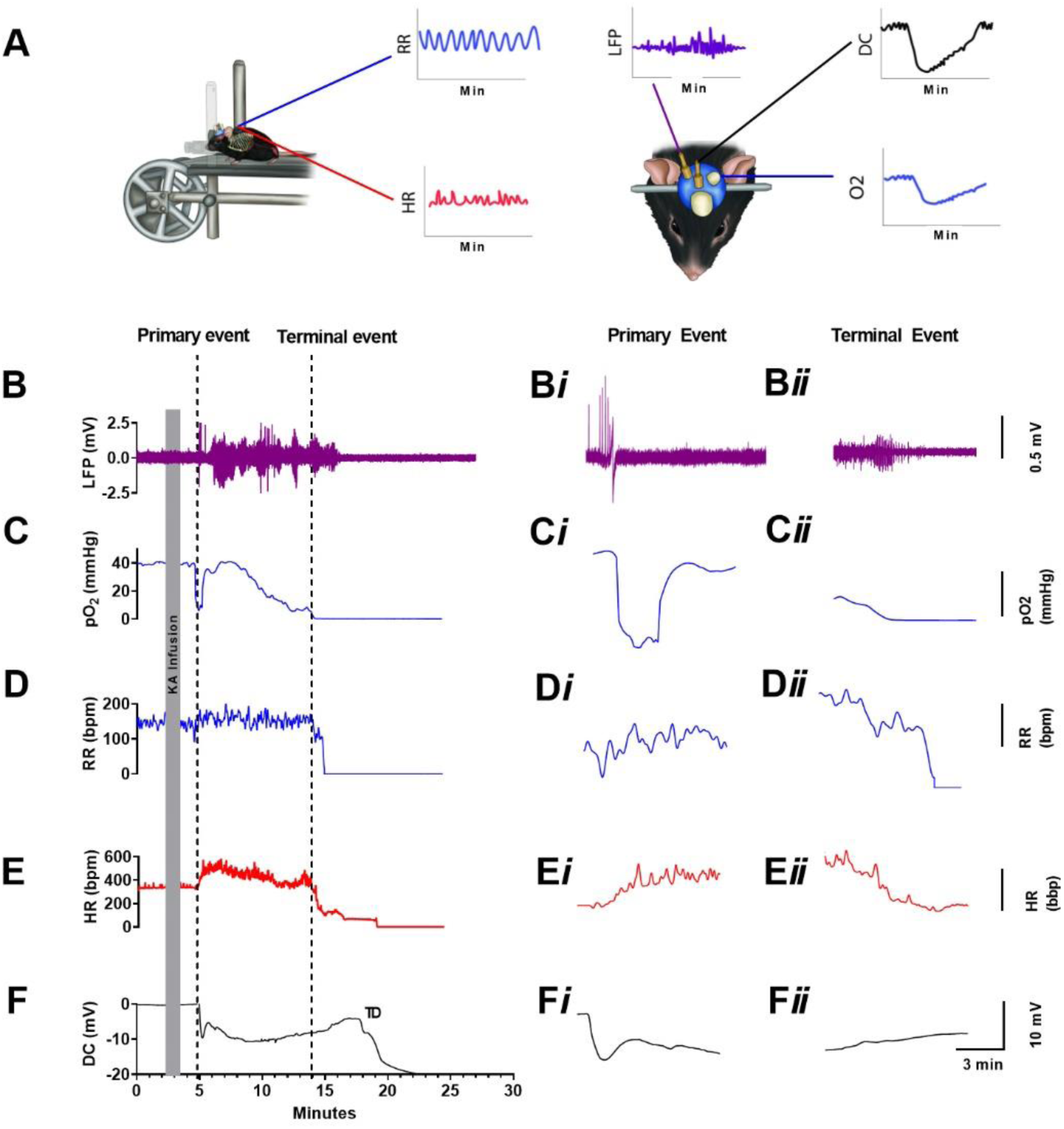
Brainstem hypoxia precedes apnea and asystole followed by spreading depolarization. Representative profile from a single C57BL/6J + kainic acid mouse. **A**. Model schematic of awake mouse experimental setup. Mice are trained on a head fixed horizontal treadmill. **B-F**. Example of kainic acid-induced seizure showing brainstem LFP, absolute oxygen, respiratory rate, heart rate, and DC spreading depolarization. Following kainic acid infusion in the HPC, a primary sequence of events occurred initiated by **B-Bi**. Epileptiform activity in the pre-Bötzinger complex (PBC) concomitant with this, there was **C-Ci**. hypoxia in the PBC, **D-Di**. unchanged respiratory rate, **E**. increase in heart rate, and finally **F-Fi**. a non-fatal DC negative shift indicating spreading depolarization. **Bii**. Fatality was initiated by a terminal seizure in the PBC, **Cii**. pO_2_ in the PBC became severely hypoxic and did not recover **Dii**. respiratory rate decreased and became terminally apneic, **Fii**. a fatal DC negative shift indicating spreading depolarization and finally **Eii**. heart rate began to decrease and became terminally asystolic.

While our acute kainic acid experiments provide proof-of-principle of the neurovascular hypothesis, to be therapeutically useful, the drugs must be on board in a chronic manner to prevent the mortality associated with self-generated seizures. Thus, we adopted the *Kcna1*^-/-^ mice that exhibit this phenotype. Given that a COX-2 blocker and LTCC antagonist both extended life in an acute model of seizure-induced premature death, the core question was whether chronic administration to *Kcna1* ^*-/-*^ mice, a more relevant model of clinical SUDEP, could also extend life.

### Chronic COX-2 inhibitors and LTCC antagonists extend life in a self-generating seizure model of SUDEP

We selected the *Kcna1*^-/-^ mouse, on a C3HeB/FeJ congenic background, because they display dramatically decreased survival compared WT littermates (Smart et al., 1998; Rho et al., 1999) and high mortality rates between postnatal day (P) 35 and P50 in our laboratory. Video and direct observation revealed that mortality was always correlated with a preceding self-generated behavioral seizure. We first exploited this high seizure and mortality epoch to assess the impact of chronic oral ibuprofen treatment. At P30, mice were randomly assigned to either the ibuprofen-in-water or water-only groups. The median survival time was significantly longer in male mice treated with ibuprofen (n=9) compared to the water only (n=9) group (P54 vs. P40, respectively; **Fig. 7A-A*i***) and female mice treated with ibuprofen (n=9) compared to the water only (n=9) group (P56 vs. P44, respectively; **Fig. 7B-B*i***). In a separate group of *Kcna1* ^-/-^ mice we also examined their seizure frequency to determine if ibuprofen had anticonvulsant effects. There was no significant effect of ibuprofen on seizure frequency indicating that the extension of life was not due to an anticonvulsant effect of ibuprofen (**Fig. 7E,F**).

**Fig. 7.**
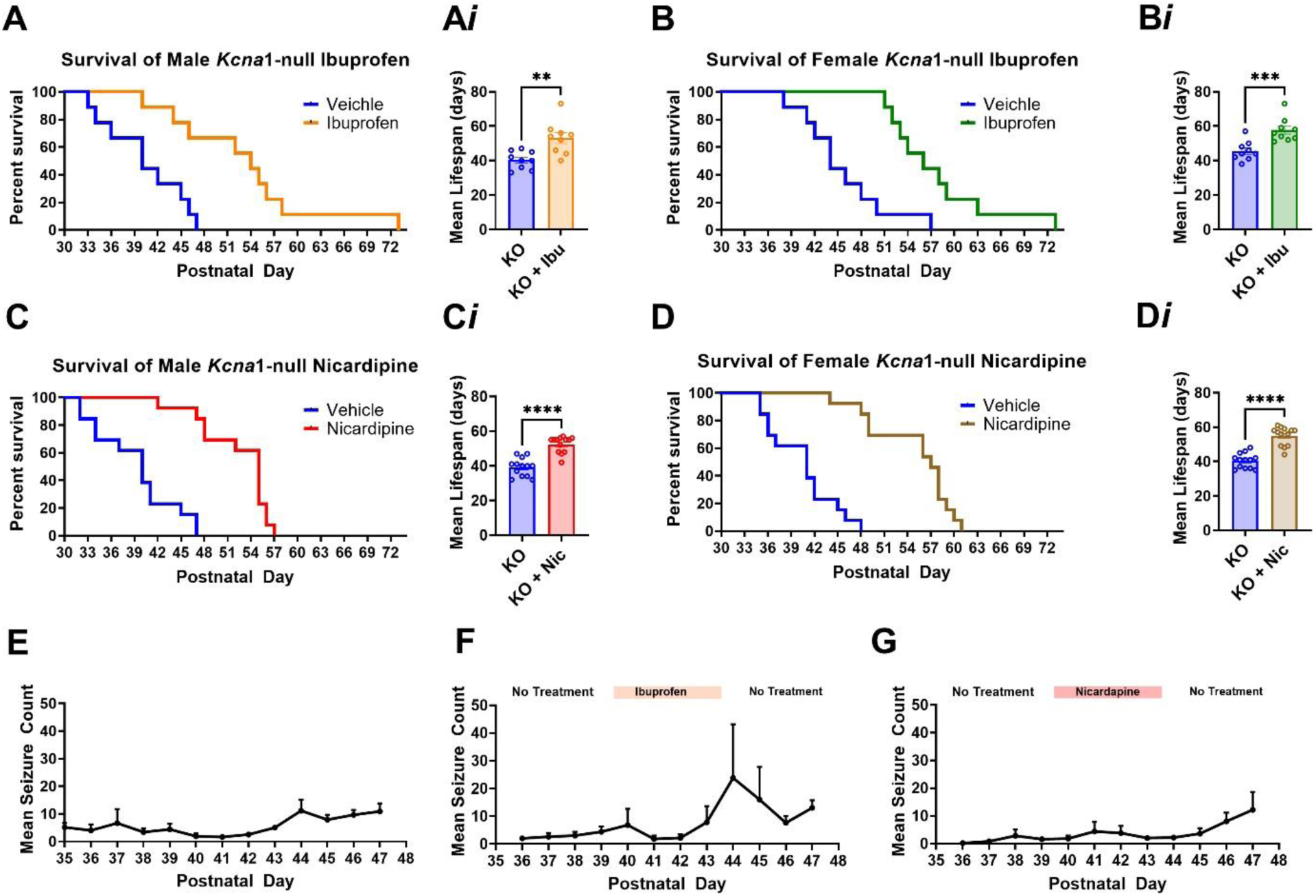
**A-B**. Chronic ibuprofen and nicardipine significantly reduces premature mortality and extends life. **A**. Male *Kcna1* ^-^/^-^ mice treated with ibuprofen lived significantly longer compared to vehicle treated mice. Median survival day for vehicle mice was 40d compared to median survival of ibuprofen of 54d **p=0.0016 (Log-rank test), **A*i***. Mean survival day for vehicle treated male mice was 40.35 ± 1.72 compared to mean survival of ibuprofen treated male mice 53.11 ± 3.20, t_16_=3.51, **p=0.0016. **B**. Female *Kcna1* ^-^/^-^ mice treated with ibuprofen lived significantly longer compared to vehicle treated mice. Median survival day for vehicle mice was 44d compared to median survival of ibuprofen treated of 56d ***p=0.0009 (Log-rank test) **B*i***. Mean survival day for vehicle treated female mice was 45.55 ± 1.87 compared to mean survival of ibuprofen treated female mice 57.66 ±2.29, t_16_=4.09, ***p=0.0009. **C-D**. Chronic Intracerebroventricular nicardipine significantly reduces premature mortality and extends life in *Kcna1* ^-^/^-^ mice. **C**. Male *Kcna1* ^-^/^-^ mice treated with nicardipine lived significantly longer compared to vehicle mice. Median survival day for vehicle mice was 40d compared to the median survival of nicardipine treated of 55d. **C*i***. Mean survival day for vehicle treated male mice was 39.23 ± 1.44 compared to mean survival of nicardipine treated male mice 52.38 ± 1.28, t_24_=6.80, ****p=<0.0001. Female *Kcna1* ^-^/^-^ mice treated with nicardipine lived significantly longer compared to vehicle treated mice. Median survival day for vehicle mice was 41d compared to median survival of nicardipine-treated mice of 57d ****p=<0.0001 (Log-rank test) **D*i***. Mean survival day for vehicle treated female mice was 40.38 ± 1.20 compared to mean survival of nicardipine treated female mice 54.84 ± 1.50, t_24_=7.52, ****p=<0.0001. Compared to controls (E), chronic administration did not affect seizure frequency in both ibuprofen (**F**) and nicardipine treated mice (**G**) (Percent change of baseline vs. treatment, ANOVA F_2,15_ = 0.3000, p=0.75.) Mean = ± SEM.

A single dose of nicardipine extended life in our acute model of seizure-induced premature mortality. To ensure dose proportionality and steady states of drug in serum, as well as avoid the peripheral side effects associated with LTCC antagonists, including severe hypotension, hypothermia, and bradycardia, we administered nicardipine intracerebroventricularly via an osmotic minipump to *Kcna1* ^-/-^ mice. The median survival time was significantly longer in male mice treated with ICV nicardipine (n=13) compared to ICV-vehicle (n=13) group (P55 vs. P40, respectively; **Fig. 7C, Ci**) and female mice treated with nicardipine compared to controls P57 vs. P41 respectively **Fig 7D, Di**. These data clearly demonstrate that chronic administration of the COX-2 blocker ibuprofen and the LTCC blocker nicardipine extends survival in a chronic model of seizure-induced premature mortality.

## Discussion

Here, we demonstrate for the first time that acute and chronic administration of the COX-2 inhibitor, ibuprofen, or the L-type calcium channel (LTCC) blocker nicardipine (which blocks postictal vasoconstriction via different mechanisms) significantly extended the life of mice in both models of seizure-induced premature mortality without reducing seizure frequency and thus provides a proof-of-principle that SUDEP has a vascular component. Our results also provide pre-clinical evidence that targeting the enzyme COX-2 or LTCC’s may be a potential therapeutic strategy for those at risk for SUDEP.

Consistent with findings found in animal studies, human neuroimaging studies have clearly delineated postictal hypoperfusion in the vicinity of the seizure onset zone and up to 90 minutes after seizure termination by means of two different imaging methodologies: arterial spin labeling (ASL)–perfusion MRI and computed tomography-perfusion (CTP) (Gaxiola-Valdez et al., 2017; Li et al., 2019; Perera et al., 2020). Moreover, Liu and colleagues studied patients with forebrain focal epilepsies using ASL MRI and found postictal hypoperfusion of medullary brainstem respiratory centres following generalized seizures providing evidence for the increased risk of SUDEP seen in bilateral tonic clonic seizures over other seizure types (Liu et al., 2020). Their study supports our neurovascular hypothesis and demonstrates that postictal hypoperfusion can occur in brainstem regions distant from the seizure onset zone. It is still unclear why a seizure becomes fatal in those who previously had similar seizures in the past. Liu and colleagues likely observed subthreshold changes in brainstem perfusion following seizures that were not severe enough to cause SUDEP. Repeated seizure activity may induce the de novo formation of medullary “lesions” and these changes become fatal during a terminal seizure (Jaster et al., 2008). Along those lines, it is plausible that repeated exposure to postictal hypoxia in the brainstem may impart an additive risk of SUDEP, only becoming fatal once a severity threshold is met. Evidence for this comes from studies showing that progressive neurodegeneration is seen in epileptic mice because of cumulative excitotoxicity and reduced regional blood flow during seizures (Meldrum, 2002; Leal-Campanario et al., 2017).

The interaction between neurovascular and neurometabolic coupling is critical to supply the high energy demands of the brain during both normal physiology and pathological conditions as seen in seizures and could possibly contribute to our neurovascular hypothesis of SUDEP (Teskey and Tran, 2021). Increased neural activity raises the metabolic rate and subsequent oxygen utilization by mitochondria which in turn reduces the availability of free oxygen in a system. The increased utilization of free oxygen coupled with the oxygen debt resulting from postictal vasoconstriction could lead to fatal outcomes if the brainstem is involved. Further, seizure activity has been clearly shown to produce reactive oxygen species (ROS) (Shekh-Ahmad et al., 2019; Olowe et al., 2020). ROS are formed when electrons are transferred to oxygen molecules which primarily occurs at the mitochondrial electron transport chain and can produce deleterious cellular damage. At low levels, ROS produce a dilatory effect on cerebral arterioles, however, increased levels produce vasoconstriction which could worsen seizure-induced postictal vasoconstriction (Carvalho et al., 2018). Further, a study by Gom and colleagues found the ketogenic diet to be protective against postictal hypoxia, raising baseline oxygen levels in a rodent study (Gom et al., 2020). Studies in both Kcna1^-/-^ and Scn1a^R1407X/+^ mice have demonstrated that the ketogenic diet significantly prolongs survival compared to mice that are fed a standard diet (Simeone et al., 2016; Teran et al., 2019). However, the mechanisms leading to this protection remain unknown. It has been postulated that the protection afforded by the ketogenic diet is likely mediated by fatty acid-induced activation of mitochondrial uncoupling proteins (Sullivan et al., 2004). These findings suggest that the ketogenic diet may be a possible treatment for those at increased risk of SUDEP.

Several rodent models have been developed in recent years to gain mechanistic insights into the pathophysiology of SUDEP. Such models include the Scn1a^*R*1407X^, RyR2^*R*176Q^, Scn1a^-/-^, Kcna1^-/-^, DBA/1&2, and Cacna1a^*S*218L^, mouse models as well as inducible kainic acid and maximal electroshock seizure models (Shen et al., 2010; Aiba et al., 2016; <u>Purnell</u> et al., 2017; Jansen et al., 2019;). One model that predominates in the literature is the DBA/2 mouse strain which is susceptible to seizure-induced respiratory arrest following an auditory tone. However, the DBA/2 model of SUDEP is subject to an important limitation, as susceptibility to seizure-induced respiratory arrest lasts only approximately 7 days (Tupal and Faingold, 2006). This temporal limitation makes it highly problematic to evaluate any chronic treatment protocol, which would be needed for human SUDEP prevention. Recurrent and uncontrolled seizures remain the principal risk factor for SUDEP. However, unlike human epilepsy, this strain does not produce self-generated seizures, reducing translatability to clinical SUDEP. Another common strain to investigate SUDEP mechanisms is the Cacna1a^*S*218L^ mouse strain, but these mice have been reported to die after a mean survival period of 2.7 days including high mortality in the days following brain electrode implant surgery (Jansen et al., 2019; Loonen et al., 2019).

Clinical case reports and retrospective studies have clearly demonstrated breathing dysfunction as a putative cause of SUDEP (Ryvlin et al., 2013). Another significant limitation in animal studies is the use of anesthetic preparations to study breathing dysfunction and seizure-induced death. Several studies use the anesthetic urethane in genetic strains (Aiba and Noebels, 2015; Aiba et al., 2016; Loonen et al., 2019). It has been established that baseline breathing and/or CO_2_ chemoreception in mice is decreased by anesthetics widely viewed as not affecting respiratory control such as urethane, and even at subtherapeutic doses (Massey and Richerson, 2017). These effects of anesthetics on breathing may alter the interpretation of studies of respiratory physiology in vivo. Additionally, a disproportionate number of cases of SUDEP occur at night and during sleep (Ryvlin et al., 2013; Sveinsson et al., 2018). It has been postulated the reason for this phenomenon, is due to the individual maintaining a prone position following a seizure, which could then lead to airway obstruction from suffocation (Tao et al., 2015). However, the specific mechanisms as to why SUDEP favours a sleep-state is still unknown. Animal studies have proposed dysfunction of CO_2_-induced arousal as a candidate cause (Buchanan, 2019). Sensitivity of the respiratory system to hypercapnia (increased blood CO_2_) and hypoxia is reduced during sleep (Santiago et al., 1984). Thus, if breathing transiently stops during a nighttime seizure, the accumulated CO_2_ may not be sufficient to mount a respiratory response to stimulate breathing due to the reduced sensitivity to CO_2_. The CO_2_ stimulus may be insufficient to stimulate arousal and restore airway patency as typically occurs with arousal. It has been established that anesthetics exert some of their effects via actions in brainstem structures involved in arousal and CO_2_ chemoreception, and therefore the use of anesthetics may blunt an arousal response to CO_2_ (Antognini et al., 2003).

Our data also indicate that brainstem SD is a consequence, rather than a cause, of seizure-induced brainstem hypoxia. Indeed, the phenomenon of SD following hypoxia was first observed in 1993 by Czeh and colleagues. They noted a SD-like event after reducing blood flow or oxygen in CA1 pyramidal cells (Czeh et al., 1993). Interestingly, they found that if blood or oxygen is restored, hypoxic SD terminates and normal function returns; if not restored, an electrical “wave of death” ensues (Zandt et al., 2011). To restore ion gradients following a hypoxic SD, oxidative energy is required (Hansen et al., 1984). During a normoxic SD, pO_2_ can decrease, however, mitochondria receive sufficient O_2_ to provide an oxidation response (Mayevsky et al., 1993; Rex et al., 1999). In contrast, in the hypoxic brain, mitochondrial enzymes become reduced, worsening the oxygen debt (Rex et al., 1999).

Contrary to our findings, a study by Jansen and colleagues found that SD caused respiratory collapse, apnea and finally hypoxia once it had invaded the ventrolateral medulla of Cacna1a^S218L^ mice (Jansen et al., 2019). Using recording electrodes and oxygen sensing probes at the level of the pre-Bötzinger complex, they found that suppression of the ventrolateral medulla following SD induces rapid apnea and subsequent hypoxia. Further, they found that fatal SD and secondary hypoxia could be reversed by resuscitation techniques. They conclude that SD was not caused by local hypoxia, but instead initiated hypoxia. The differences seen between both our study and Jansen’s (2019) could be attributed to differences in seizure networks between SUDEP animal models, as well as inherent strain differences. In fact, Jansen and colleagues observed reduced survival in their mice due to an increased risk of SD following tissue trauma from electrode implantation.

Sudden unexpected death in epilepsy continues to be a major cause of death in people with drug resistant epilepsy (Holst et al.,2013; Massey et al., 2014). Currently, no diagnostic test or drug treatment exists for SUDEP, and no specific post-mortem findings have been identified as pathognomonic for SUDEP. Our data indicate that administration of the COX-2 inhibitor ibuprofen, or the LTCC blocker nicardipine significantly extends life in two models of seizure-induced premature mortality, revealing a promising strategy for future therapy in people at risk of SUDEP. COX-2 inhibitors are readily available over the counter and prescription NSAIDs are also used in the reduction of inflammation after injury (Ong et al., 2007). Nicardipine is prescribed for the management of hypertension and used in emergency cases for subarachnoid hemorrhage and cerebral vasospasm (Narotam et al., 2008). Clinical trials are warranted to investigate the potential therapeutic efficacy of these or similar medications in reducing SUDEP risk.

## Materials and Methods

### Mice

Adult male and female C57BL/6J (Jackson Laboratory, JAX: 000664) and *Kcna1* ^*-/-*^ (in-house breeding) mice on a C3HeB/FeJ congenic background were used (aged 7-10 weeks and weighing between 23–31 g). Mice were housed individually in clear plastic cages and were maintained on a 12:12 hr light/dark cycle with lights on at 07:00 hr, in separate colony rooms under specified pathogen free conditions. Food and water were available *ad libitum* and mice were allocated randomly to different test groups. Mice were handled and maintained according to the Canadian Council for Animal Care guidelines. All procedures were approved by the Health Sciences Animal Care Committee at the University of Calgary (AC16-0272, AC17-0191).

### Surgery

#### Brain implantation surgery

Mice were anesthetized with 5% isoflurane and maintained on 2% isoflurane mixed with 100% oxygen. The skull was secured in a stereotaxic apparatus (David Kopf Instruments) using ear and incisor bars. Diluted lidocaine (0.5% 1:4 lidocaine, saline) was injected subcutaneously over the incision site followed by a sagittal incision along the midline of the head. Burr holes were drilled in the skull at stereotaxic coordinates. Bipolar depth electrodes were implanted unilaterally into the dorsal hippocampus at these coordinates relative to bregma: (AP) -2.2, (ML)-2.5, (DV) -2.0 and the pre-Bötzinger complex at: (AP) -7.0, (ML) -1.4, (DV) -4.1. Oxygen sensing optodes (Oyxylite, Oxford Optronix) were implanted in the ipsilateral ventral hippocampus at: (AP) -1.5, (ML) -0.8, (DV) -1.5 and in the contralateral pre-Bötzinger complex at: (AP) -7.0, (ML) +1.4, (DV) 4.1. A 22-gauge infusion guide cannula (Plastics One, Roanoke, VA) was implanted in the contralateral dorsal hippocampus at: (AP) -2.7, (ML) +3.0, (DV) -3.0. For the three-structure experiment, an optode was also placed in the nucleus of the solitary tract (NTS) at these coordinates relative to bregma: (AP) -6.38, (ML) +0.4, (DV) -7.1. Electrodes and probes were secured to the skull using Metabond Quick Adhesive (C&B) and dental acrylic. Mice were administered injectable buprenorphine (0.05 mg/kg) every 12 hours for 3 days post-operatively and mice recovered for 3 days before diaphragmatic surgery.

#### Diaphragmatic surgery

Mice were anesthetized with 5% isoflurane and maintained on 2% isoflurane mixed with 100% oxygen. Diluted lidocaine (0.5% 1:4 lidocaine, saline) was injected subcutaneously at the incision site on the dorsal segment of the ribs just below the ribcage. Electrode wires were implanted as described by Pagliardini et al., (2012). In brief, multi-stranded Teflon-coated stainless-steel electrode wire (#AS633, Cooner Wire) were passed through the diaphragm using a suture needle and secured in place by a knot. A second electrode wire was placed 1 mm away from the first to provide a bipolar recording. The electrode ends were then tunneled under the skin to an incision on the animal’s dorsum, between the scapulae. The uninsulated ends of the electrode wires were then attached to male amphenol pins and secured in a 9-pin ABS socket (Ginder Scientific) that protruded from the back. The incision sites were sutured, and the mice recovered. Mice were administered injectable buprenorphine (0.05 mg/kg) every 12 hours for 3 days post-operatively and were given 7 days to recover before beginning experiments.

#### Electrocardiography surgery

Mice were anesthetized with 5% isoflurane and maintained on 2% isoflurane mixed with 100% oxygen. Diluted lidocaine (0.5% 1:4 lidocaine, saline) was injected subcutaneously at the incision site on the ventral aspect of the mouse from below the neck down to the caudal abdomen. A midline incision was made exposing underlying musculature. Multi-stranded Teflon-coated stainless-steel electrode wire (#AS633, Cooner Wire) were placed in the xiphoid cartilage, the right forelimb in the cleidocephalicus muscle and the left hindlimb in the rectus femoris-quadricep muscle (Hedrich 2004). All electrodes were tacked to musculature using suture. The electrodes were then tunneled under the skin and exited through a mid-scapular incision. The uninsulated ends of the electrode wires were then attached to male amphenol pins and secured in a 9-pin ABS socket (Ginder Scientific) that protruded from the back. The incision sites were sutured, and mice were recovered from anaesthesia. Mice were administered injectable buprenorphine (0.05 mg/kg) every 12 hours for 3 days post-operatively and were given 7 days to recover before beginning experiments.

#### Intracerebroventricular surgery

On postnatal day 29, mice were anesthetized with 5% isoflurane and maintained on 2% isoflurane mixed with 100% oxygen. The skull was secured in a stereotaxic apparatus using ear and incisor bars. Diluted lidocaine (0.5% 1:4 lidocaine, saline) was injected subcutaneously over the incision site followed by a sagittal incision along the midline of the head. The incision was then extended over the dorsum of the mouse posterior to the scapulae. Haemostats were introduced subcutaneously to widen incision and produce dead space for pump insertion. A burr hole was made over the right lateral ventricle at these coordinates relative to bregma: (AP) + 0.3, (ML) -1.0, (DV) -3.0 (DeVos & Miller, 2013). The infusion cannula was secured to the skull using Metabond Quick Adhesive (C&B) and dental acrylic. Mice were administered injectable buprenorphine (0.05 mg/kg) every 12 hours for 1 day post-operatively and mice were allowed to recover for 1 day before starting the experiment on P30. The patency and placement of the cannula was verified at the end of the experiments using methylene blue dye (Sigma-Aldrich) administration.

### Histology

At the end of each experiment, brains were harvested and stored in 4% paraformaldehyde for a minimum of 24 hrs followed by sucrose. Coronal sections (40 μm) were collected using a freezing sliding microtome. Slices of interest were stained with cresyl violet dye (Sigma-Aldrich) and mounted on glass slides to identify damage tracts to ensure correct placement of electrodes, cannulas and probes using light microscopy.

### Acute Experiments

#### Respiratory dysfunction and failure precede asystole

Male C57BL/6J and *Kcna1* ^-^/^-^ were chronically implanted with bipolar electrodes in the hippocampus and pre-Bötzinger to record LFP, and oxygen sensing optrodes to measure absolute oxygen. Bipolar diaphragmatic EMG electrodes obtained continuous cardiac and respiratory rate. Kainic acid (1.4 μg in 0.4 μL) was infused through an intrahippocampal cannula and animals were recorded until death. Eight C57BL/6J and eight *Kcna1*^*-/-*^ mice were used to derive representative date.

#### Acute dosing of ibuprofen and nicardipine

Male and female C57BL/6J mice (Jackson Laboratory, JAX: 000664) were administered either 15, 50 or 100 mg/kg ibuprofen (Cayman Chemicals) or vehicle (DMSO, Sigma-Aldrich) intraperitoneally (i.p.) 30 minutes prior to infusion of kainic acid through an intrahippocampal cannula. Kainic acid (1.4 μg in 0.4 μL in males and 2.8 μg in 0.8 μL in females) was infused unilaterally into the right dorsal hippocampus at 0.1 μL/min through a 1 μL Hamilton syringe (Hamilton Robotics, Reno NV) and micro-syringe pump (Harvard Apparatus, model 55-2222, Canada). In the repeated dosage of ibuprofen experiments, male C57BL/6J mice were administered an initial loading dose of 100 mg/kg ibuprofen or vehicle i.p. 30 minutes prior to infusion of kainic acid, followed by 3 successive maintenance doses of 50 mg/kg, every 2 hours. In the LTCC blocker experiments, male and female C57BL/6J mice were administered 5 mg/kg nicardipine (Caymen Chemicals) or vehicle (DMSO) 5 minutes prior to infusion of kainic acid. In all experiment’s LFP, EMG, and brain pO_2_ were recorded continuously and survival time following infusion was recorded in each animal.

#### Ketamine, diazepam, isoflurane

Male C57BL/6J mice were administered 100 mg/kg ibuprofen i.p. 30 minutes prior to infusion of kainic acid. Two hours post kainic acid infusion, 100 mg/kg ketamine (Vetoquinol) was given i.p. This dose produced marked to moderate anesthesia, therefore animals were placed on a heating pad to maintain normothermia. In the diazepam experiments, male C57BL/6J mice were administered 100mg/kg ibuprofen i.p. 30 minutes prior to kainic acid infusion. Two hours post kainic acid infusion, 5 mg/kg diazepam (Sandoz) was given i.p. In the isoflurane experiments, male C57BL/6J mice were administered 100 mg/kg ibuprofen i.p. prior to kainic acid infusion. After 2 hours, mice were anesthetized with 4% isoflurane (Baxter) mixed with 100% oxygen and titrated to 2% isoflurane for 2-hours. In the ketamine and diazepam experiments, mice were administered 100 mg/kg ketamine and 5 mg/kg diazepam after kainic acid infusion. In the diazepam and isoflurane experiments, mice were given 5 mg/kg diazepam and 4% isoflurane titrated to 2%. In all experiments, LFP, EMG and brain pO_2_ were recorded for 8-hours post kainic acid infusion. After the 8-hours, mice were transferred to their home cages for observation.

#### Three brain structure analysis: HPC, PBC, and NTS

Male C57BL/6J mice were chronically implanted with oxygen sensing optrodes in the hippocampus, pre-Bötzinger and nucleus of the solitary tract. Kainic acid (1.4 μg in 0.4 μL) was infused through an intrahippocampal cannula and animals were recorded until death. For better temporal analysis, the sample rate was increased from the standard 20 pulses/min (0.333 Hz) to 60 light pulses/ min (1 Hz).

### Chronic Experiments

#### Ibuprofen in drinking water

Male and female *Kcna1* ^*-/-*^ mice on a C3HeB/FeJ congenic background were used. These mice carry a mutation of the *Kcna1* gene on chromosome 6 as a result of gene targeted deletion. Starting the experiment on postnatal day 30, male and female mice received either water or water with 0.6 mg/mL ibuprofen sodium (Sigma-Aldrich). Mice were monitored every 12 hours and date of death recorded. Ibuprofen was protected from light and replaced every 12 hours to ensure photostability and solubility.

#### Intracerebroventricular nicardipine administration

Male *Kcna1* ^-/-^ were implanted with Alzet 42-day mini-pumps (model 2006; Durect, Cupertino) which continuously delivered nicardipine into the left lateral ventricle (2 mg/mL, pumping rate 0.15 μL/hr) this dose was selected based on pilot data suggesting toxicity and lethality at higher concentrations. Nicardipine was dissolved in DMSO, PEG and 100% ETOH (50:40:10). Mice were monitored every 12 hours and date of death recorded.

#### Anti-seizure drug screening of ibuprofen and nicardipine

To assess whether ibuprofen and/or nicardipine exerted antiseizure effects we analyzed seizure frequency, utilizing video-cortical LFP monitoring. *Kcna1*^*-/-*^ mice, at P28, were implanted with stainless steel screw electrodes to record LFP (Pinnacle Technology Inc., #8247) at the following coordinates relative to bregma: +1.0mm (AP) and ±1.0mm (ML), and -2.00mm (AP) and ±2.00mm ML), and -5.0mm (AP) and 0.0mm (ML). Mice were allowed one week of recovery after which they were connected to the acquisition system (Pinnacle Technology Inc., #8200-K1-SE3) and recordings commenced. Video-EEG data was acquired continuously for 12 days at 250 Hz and 30 frames per second. Seizures were quantified using a custom, semi-automated seizure detection algorithm (MATLAB, MathWorks). Detected seizures were then verified visually and assigned a Racine score by a blinded and experienced scorer. Mice were recorded continuously 24 hours per day for 12 days. Starting on P36 a 3-day baseline was recorded, followed by 6 days of either ibuprofen or nicardipine in water, ending the recordings with a 3 day washout period.

### Absolute oxygen recordings

Absolute oxygen recordings were measured using an implantable fibre-optic oxygen-sensing probe (Oxylite, Oxford Optronics). These probes emit a short burst of LED light to a platinum fluorophore at the tip of the probe. When oxygen molecules collide with the fluorophore the light emitted by the probe is quenched and is registered as a decay in luminescence which is relayed to the terminal for display. The probe allows for accurate and continuous readings of oxygen in brain parenchyma in the awake and freely moving animal. The optode can measure a volume of 500 μm^3^ between 0 and 90 mmHg. Unless otherwise stated, sample rate of 60 light pulses/ min were used at 1 Hz. Severe hypoxia is defined as <10mmHg because oxygen levels at or below this threshold are associated with brain injury resulting from cellular damage and death leading to worse clinical outcomes (Oddo et al., 2011).

### Electroencephalography recordings and analysis

EEG was recorded using a Grass Neurodata Acquisition System (Model 12C, Quincy, MA) with digital EEG-amplifier. The recorded EEG analog signal was digitized with DATAQ software (DATAQ Instruments, Akron, OH). Behavioral seizures were scored using the Racine scale: Stage 0: freezing accompanied by an electrographic seizure; Stage 1: motor arrest accompanied by facial automatisms; Stage 2: head nodding and chewing; Stage 3: unilateral forelimb clonus; Stage 4: rearing on hind limbs with bilateral forelimb clonus; Stage 5: rearing, loss of posture, and clonus of all four limbs (Racine, 1972).

### Spreading depolarization recordings and analysis

LFP’s in the brainstem were recorded with an unipolar electrode and amplified with a DC amplifier (MultiClamp 700B) and digitized at 200Hz (Digidata 1440A) The DC aspect of this signal was extracted using a 1 dimensional gaussian filter (scipy.ndimage.gaussian_filter1d, sigma=1s) (Harris et al., 2020). LFP oscillations riding on this signal were extracted by subtracting the raw signal by the filtered DC signal (low-pass 100Hz).

### Electromyography recordings and analysis

Heart rate and respiratory rate was obtained from raw EEG signal as a post-processing procedure. To obtain respiratory rate, the signal was low pass filtered below 0.01 Hz using a fifth-order Butterworth filter to remove signal drift and to determine during which part of the recordings the mouse was alive. Next, the raw EMG data was again low pass filtered below 4Hz to extract respiratory information. The duration of the recording while the mouse was alive was the analyzed for individual breaths, done by applying a 1D data peak detection algorithm (peakutils.peak). Local maxima larger than a normalized threshold height were identified as breaths (the exact amplitude being dependent on the signal strength in the individual dataset). Heart rate data was processed by using custom algorithm (Python 2.7.12, scipy 0.18.1, peakutils 1.1.0). The signal was low pass filtered at 5Hz using a fifth-order Butterworth filter to extract respiration and movement artifact. This low frequency signal was subtracted from the raw EMG to remove low frequency components, leaving only high frequency components such as heartbeats and signal noise. Individual heartbeats were determined by identifying local maxima using a 1D data peak detection algorithm (peakutils.peak). Only peaks over a certain threshold amplitude defined relative to the maximum amplitude in the signal (the exact threshold being dependent on the overall signal strength in the individual dataset) were accepted as heartbeats. A minimum distance between peaks was defined as 12ms for heartbeats, as heartbeats at a higher frequency would not be physiologically possible. Breathing and heart rate were obtained by calculating the time difference between peaks and creating a moving average across the entire data set.

### Statistics

All statistical analyses were performed using Prism version 9.3.1 (GraphPad, La Jolla, CA). Unpaired (comparisons between animals) t-tests were used for experiments with only two groups. When more than two groups were compared either a one-way ANOVA, or two-way ANOVA were applied. Survival curves were analyzed by Kaplan–Meier survival estimates using the log-rank test. All data were tested for normality before additional statistical testing (i.e., post-hoc analysis). Significance thresholds were set to: *p<0.05. Unless otherwise stated, values were expressed as means ± standard error of the mean (SEM). Determination of sample and effect size was based on approximations informed by pilot study data from our laboratory. Further, our foundational work (Farrell et al., 2016) reflected the sample size of our current study, showing the minimum number of animals required to derive statistically significant oxygen data. The data in all figure panels reflect multiple experiments performed on different days or representative traces of multiple animals. In this study, we used a minimum of 4 mice per group (i.e., biological replicates) for all experiments or a maximum of 9 mice per group. Statistical details, including sample size, are found in the results section. All outliers were included in summary figures and used in statistical analysis. Our dependent variable was time to death; therefore animals were used once per group/experiment.

## Data Availability

All source data files for summary figures have been deposited with Zenodo at the following URL: https://zenodo.org/record/6369179#.YjY3herMKUk. The source code files for the seizure detection algorithm are accessible at Zenodo under the following DOI: 10.5281/zenodo.6369179.

## Acknowledgements

Canadian Institutes of Health Research Grant (PJT-152956) to G.C.T. and SUDEP Aware to A.G.G. J.S.F. was funded by a CIHR postdoctoral fellowship.

---

**Suppl Fig. 1.**
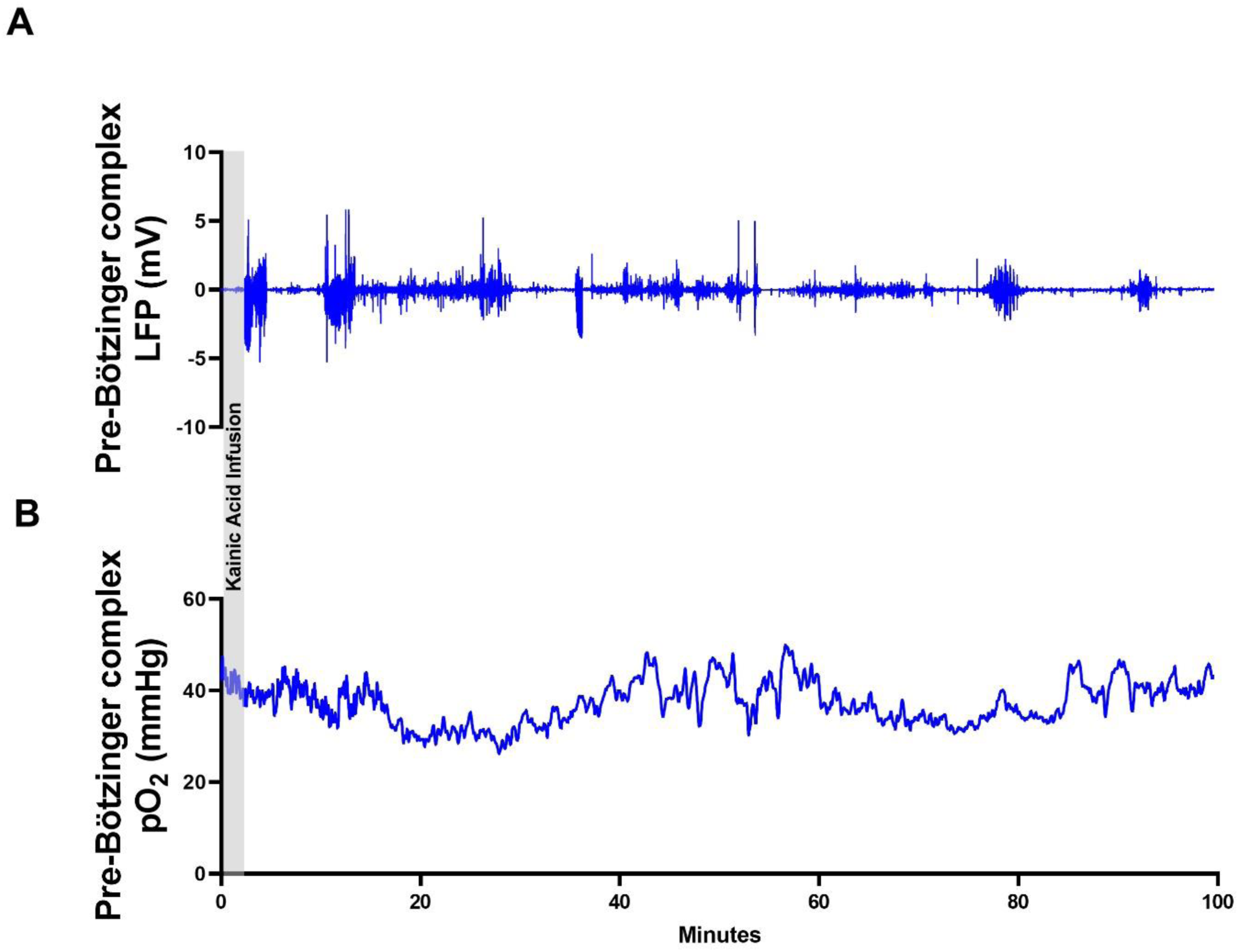
Acute administration of ibuprofen protects the pre-Bötzinger complex from seizure-induced hypoxia. Representative profile from a C57BL/6J + kainic acid mouse injected with 100mg/kg ibuprofen 30 minutes prior to kainic acid infusion (no differences were observed between C57BL/6J and *Kcna1*^-/-^ mice). **A**. Mice treated with ibuprofen displayed electrographic seizure around the same time vehicle mice died, however, **B**. brainstem pO_2_ remained within normoxic range.

## Notes

### Competing Interest Statement

The authors have declared no competing interest.

